# Structural basis for the inhibition of coronaviral main proteases by PF-00835231

**DOI:** 10.1101/2024.04.22.590578

**Authors:** Xuelan Zhou, Xiaolu Lu, Cheng Lin, Xiaofang Zou, Wenwen Li, Xiangyi Zeng, Jie Wang, Pei Zeng, Weiwei Wang, Jin Zhang, Haihai Jiang, Jian Li

## Abstract

The main protease (M^pro^) of coronaviruses plays a key role in viral replication, thus serving as a hot target for drug design. It has been proven that PF-00835231 is promising inhibitor of SARS-CoV-2 M^pro^. Here, we report the inhibition potency of PF-00835231 against SARS-CoV-2 M^pro^ and seven M^pro^ mutants (G15S, M49I, Y54C, K90R, P132H, S46F, and V186F) from SARS-CoV-2 variants. The results confirm that PF-00835231 has broad-spectrum inhibition against various coronaviral M^pro^s. In addition, the crystal structures of SARS-CoV-2 M^pro^, SARS-CoV M^pro^, MERS-CoV M^pro^, and seven SARS-CoV-2 M^pro^ mutants (G15S, M49I, Y54C, K90R, P132H, S46F, and V186F) in complex with PF-00835231 are solved. A detailed analysis of these structures reveal key determinants essential for inhibition and elucidates the binding modes of different coronaviral M^pro^s. Given the importance of the main protease for the treatment of coronaviral infection, structural insights into the M^pro^ inhibition by PF-00835231 can accelerate the design of novel antivirals with broad-spectrum efficacy against different human coronaviruses.

## Introduction

In late 2019, a novel coronavirus disease caused by SARS-CoV-2 erupted in Wuhan, China [1–3]. SARS-CoV-2 belongs to the β-Coronaviridae family, which is in the same family as Middle East Respiratory Syndrome Coronavirus (MERS-CoV) and SARS-CoV. All three strains are highly pathogenic [4,5]. However, SARS-CoV-2 mutates easily, and the emerged variants have raised concerns about the characteristics of the virus, including transmissibility and antigenicity. WHO had identified five variants as the variants of concern (VOCs), namely B.1.1.7 (Alpha, α), B.1.351 (Beta, β), P.1 (Gamma, γ), B.1.617.2 (Delta, δ), B.1.1.529 (Omicron), and several variants as the variants of interest (VOI), including C.37 (Lambda, λ) (https://www.who.int/activities/tracking-SARS-CoV-2-variants). The spread of SARS-CoV-2 as well as its variants has lasted for more than four years and has caused more than 774 million cases of COVID-19 as of 11 February 2024, of which 7.03 million have died (https://covid19.who.int/). Many health agencies are looking for treatment options, and many drugs used for treating SARS-CoV-2 infection are also in clinical development [6–12]. One such strategy is targeting main protease (M^pro^), to selectively inhibit coronaviral replication [13–15].

The coronaviral main protease is also known as the 3C-like protease (3CLpro) [14,15]. After successfully infecting the host cells, the viral genome encodes two large overlapping polyproteins, namely pp1a and pp1ab. M^pro^ is able to process the polyproteins to produce several nonstructural proteins (NSPs) necessary for viral replication. Additionally, M^pro^ is highly conserved among β coronaviruses [15]. The recognition site of coronaviral M^pro^ depends on Gln at the P1 position, and no proteases in humans share similar cleavage site [13,14]. Therefore, M^pro^ of coronaviruses plays an important role in viral replication and the selected M^pro^ inhibitors should have broad-spectrum properties.

To date, various inhibitors targeting coronaviral M^pro^ have been developed by using drug discovery strategies such as high-throughput screening, structure-based drug design, and drug re-purposing [8,11,12,14]. Among these, PF-07321332 and PF-00835231 represent state-of-the-art inhibitors with therapeutic potential [8,11,16]. PF-07321332 share structural similarity with GC376 and has been approved for clinic application in combination with ritonavir [8,17–19]. PF-00835231 is an inhibitor designed for the main protease of SARS-CoV emerged in 2003, but its clinical trials were suspended due to the rapid disappearance of the SARS outbreak [16]. Based on the high M^pro^ similarity (96%) between SARS-CoV-2 and SARS-CoV, PF-00835231 can be a promising drug candidate for the treatment of SARS-CoV-2 infection [16]. Previous data have shown that PF-00835231 has a good inhibitory effect against SARS-CoV-2, but its oral bio-availability is relatively poor [20]. In this regard, there are two ways to increase its bio-availability. One way is to convert PF-00835231 into its phosphate pro-drug form (PF-07304814) as an intravenous treatment option, and another way is to optimize the PF-00835231 structure, largely based on the molecular basis of PF-00835231 in inhibiting various coronaviral M^pro^s.

In this study, we used a structure-based approach to investigate the inhibition efficacy and molecular basis for M^pro^ inhibition by PF-00835231. We found that PF-00835231 broadly inhibited SARS-CoV-2 M^pro^ and its mutants (G15S, M49I, Y54C, K90R, P132H, S46F and V186F). These mutations can be found in different SARS-CoV-2 variants: B.1.351 Beta (K90R), C.37 Lambda (G15S), Delta AY.4 (Y54C), BA.5 Omicron (M49I), B.1.1.529 Omicron (P132H), B.1.1.529 Omicron (V186F), and B.1.1.529 Omicron (S46F). The crystal structures of M^pro^s from SARS-CoV-2, SARS-CoV-2 VOC/VOIs, SARS-CoV and MERS-CoV M^pro^s bound to PF-00835231 were solved, and revealed the structural similarities and differences of PF-00835231 in binding different M^pro^s. The results provide structural insights into understanding the precise inhibition mechanism of this inhibitor against different M^pro^s and strongly suggest that PF-00835231 has the potential to develop broad-spectrum drug candidates. Furthermore, this study provides critical information for the optimization of PF-00835231 and the design of more effective anti-coronaviral inhibitors.

## Materials and Methods

### Expression and Purification of coronaviral M^pro^s

According to the previous experimental methods [18,21], the genes encoding SARS-CoV-2, SARS-CoV, and MERS-CoV M^pro^s were inserted into the pET-28a vector and then the recombinant plasmids were constructed. The plasmid carrying SARS-CoV-2 M^pro^ gene was used as a template for site-directed mutagenesis to generate a variety of SARS-CoV-2 M^pro^ mutants, including G15S, S46F, M49I, Y54C, K90R, P132H, and V186F. Then the recombinant plasmids were introduced into competent *Escherichia coli* Rosetta DE3 cells for protein expression. The expression and purification of wild type M^pro^ and M^pro^ mutants were carried out according to the methods described in the previous article [18,21]. Further, TEV protease was used to remove the N-terminal His tag.

### Enzymatic Inhibition Assay

Fluorogenic substrates as a donor and quencher pair were commercially synthesized. The inhibiting efficacies of PF-00835231 against different coronaviral M^pro^ proteins were detected using the fluorescence resonance energy transfer (FRET)-based enzymatic assays which has been reported previously [22]. Briefly, PF-00835231 was dissolved in DMSO to prepare a stock solution of 10 mM in advance. Then PF-00835231 was subjected to a 3-fold serial dilution in triplicate and incubated with different coronaviral M^pro^ proteins and SARS-CoV-2 M^pro^ mutants for 30 min. Subsequently, a FRET substrate was added to the reaction system, followed by another 20 min of incubation. The system was monitored and the fluorescence was recorded during the reaction. Using the GraphPad Prism software, inhibition activities (%) of PF-00835231 against coronaviral M^pro^s were finally determined.

### Crystallization

The recombinant M^pro^ proteins were concentrated to 10 mg/mL, and incubated on ice with PF-00835231 at 1:5 molar ratio for 30 minutes. Crystallization was carried out at 18_ by using hanging drop vapor-diffusion method. After 3 to 5 days, the crystals of M^pro^s in complex with PF-00835231 were obtained. The final crystallization condition of SARS-CoV-2 M^pro^-PF-00835231 complex was 0.1 M HEPES sodium pH 7.5, 10% v/v 2-Propanol, 20% w/v PEG 4000. The final crystallization condition of SARS-CoV M^pro^-PF-00835231 complex was 0.1 M HEPES pH 7.5, 12% w/v PEG 8000, 10% w/v ethylene glycol. The final crystallization condition of MERS-CoV-M^pro^-PF-00835231 complex was 0.2 M Sodium chloride, 20% w/v PEG 3350. The final crystallization condition of SARS-CoV-2 M^pro^ (Y54C)-PF-00835231 complex was 0.7 M Sodium citrate tribasic dihydrate, 0.1 M Bis-Tris propane pH 7.0. The final crystallization conditions of other SARS-CoV-2 M^pro^ mutants (including G15S, S46F, M49I, K90R, P132H, and V186F) in complex with PF-00835231 were 0.1 M-0.25 M Na_2_SO_4_, 20% w/v to 25% w/v PEG3350.

### Data Collection, Structure Determination, and Refinement

Before collecting the data, the crystals were soaked in a cryoprotective solution containing 20% glycerol and then stored in liquid nitrogen. Diffraction data were collected at 100 K on the macromolecular crystallographic beamline 02U1 (BL02U1) and 10U2 (BL10U2) at the Shanghai Synchrotron Radiation Facility (SSRF, Shanghai, China). All data sets were processed using HKL2000 software [23]. The phase problem was solved by molecular replacement. Structures were refined for several cycles in Phenix in order to achieve the desired resolution [24].

## Results

### Broad-spectrum inhibitory activity of PF-00835231

The SARS-CoV-2 M^pro^ and M^pro^ mutants (G15S, M49I, Y54C, K90R, P132H, S46F and V186F) variants were expressed and purified as previously reported [18,21]. The inhibitory activities of PF-00835231 against M^pro^s were detected by using fluorescence resonance energy transfer (FRET) assay. The results showed that PF-00835231 can effectively inhibit SARS-CoV-2 M^pro^ and its mutants. The IC_50_ value of PF-00835231 against SARS-CoV-2 M^pro^ was 0.0086µM, while the IC_50_ values of PF-00835231 against SARS-CoV-2 M^pro^ mutants were ranging from 1.2 to 3.7 µM (Figure 1). Based on our previous work, PF-00835231 also inhibited M^pro^s of HCoV-NL63, HCoV-HUK1, MERS-CoV, and SARS-CoV [25]. The Ki values of PF-00835231 against M^pro^s from various coronaviruses including HCoV-NL63, HCoV-229E, PEDV, FIPV, HKU4-CoV, HCoV-OC43, and HCoV-HKU1 range from 30 pM to 4 nM [11]. In addition, PF-00835231 exerts antiviral activity against SARS-CoV and HCoV-229E with very low semi-maximum effective concentration (EC_50_) values [16]. These data suggest that PF-00835231 has a broad enzymatic inhibitory effect against coronavirus M^pro^s, as well as this compound’s potential in preventing the current and next coronavirus pandemic.

**Figure 1.**
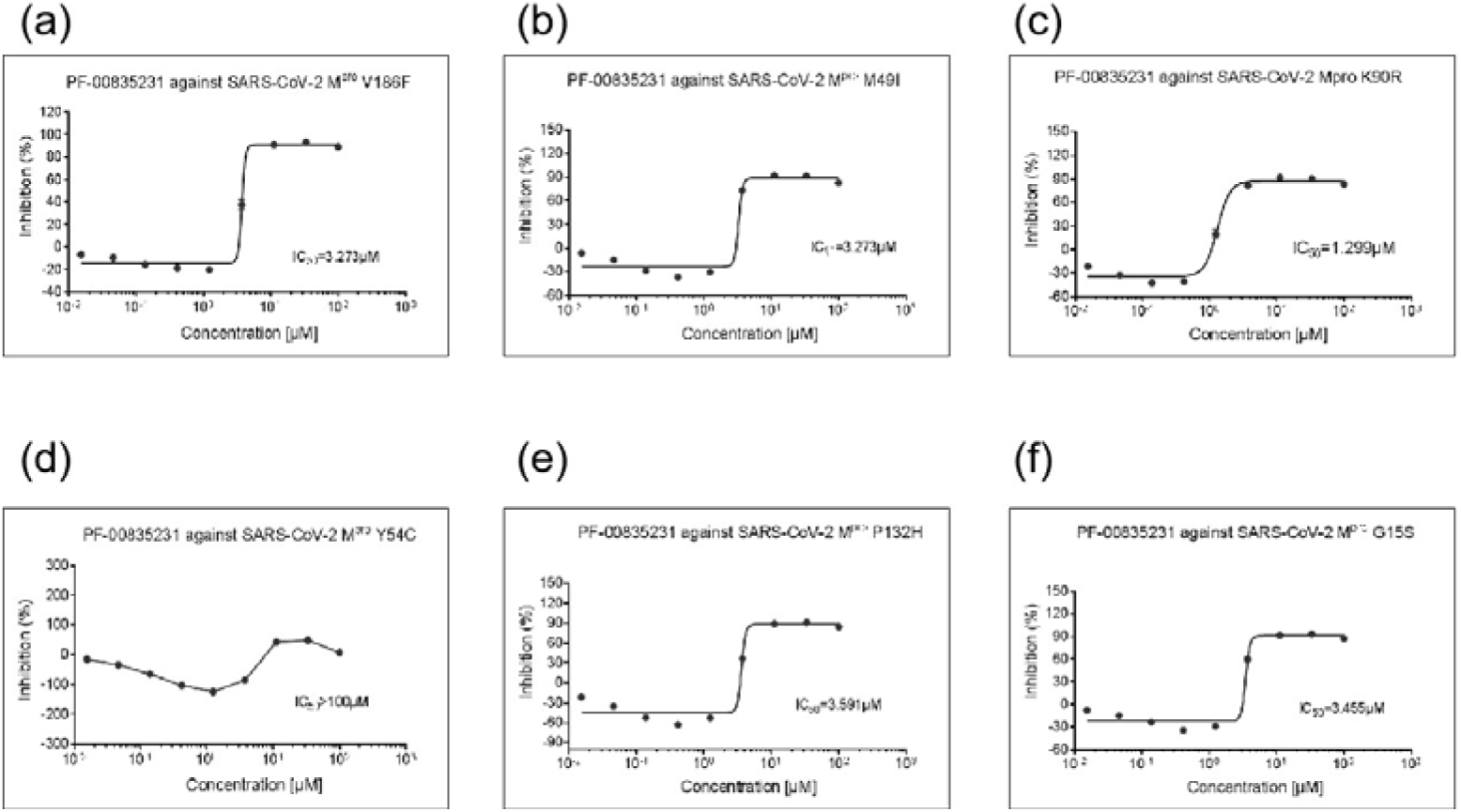
Enzymatic inhibition of PF-00835231 against SARS-CoV-2 M^pro^ and SARS-CoV-2 M^pro^ mutants. (a) Inhibition potency of PF-00835231 against the main protease of SARS-CoV-2 M^pro^ V186F. (b) Inhibition of PF-00835231 against the main protease of SARS-CoV-2 M^pro^ M49I. (c) Inhibition of PF-008352’31 against the main protease of SARS-CoV-2 M^pro^ K90R. (d) Inhibition of PF-00835231 against the main protease of SARS-CoV-2 M^pro^ Y54C. (e) Inhibition of PF-00835231 against the main protease of SARS-CoV-2 M^pro^ P132H. (f) Inhibition of PF-00835231 against the main protease of SARS-CoV-2 M^pro^ G15S.

### Crystal structure of SARS-CoV-2 M^pro^ in complex with PF-00835231

In order to understand the inhibition mechanism of PF-00835231 against SARS-CoV-2 M^pro^, we solved the crystal structure of SARS-CoV-2 M^pro^ in complex with PF-00835231. The resolution of the complex structure is 2.21Å. Data collection and refinement statistics are summarized in Table 1. As shown in Figure 2A, SARS-CoV-2 M^pro^ displays as a homodimer in the complex structure that can be divided into three subdomains, namely domain I (residues 3 to 99), domain II (residues 100 to 199), and domain III (residues 201 to 300). Domains II and III are connected by a long ring (residue 175 to 200). The narrow cavity between domain I and domain II contains the inhibitor PF-00835231, which is present in both protomer A and protomer B. An enlarged view of the narrow cavity revealed that PF-00835231 occupied the S1, S1’, and S2 subsites of SARS-CoV-2 M^pro^ in an extended conformation (Figure 2b).

**Figure 2.**
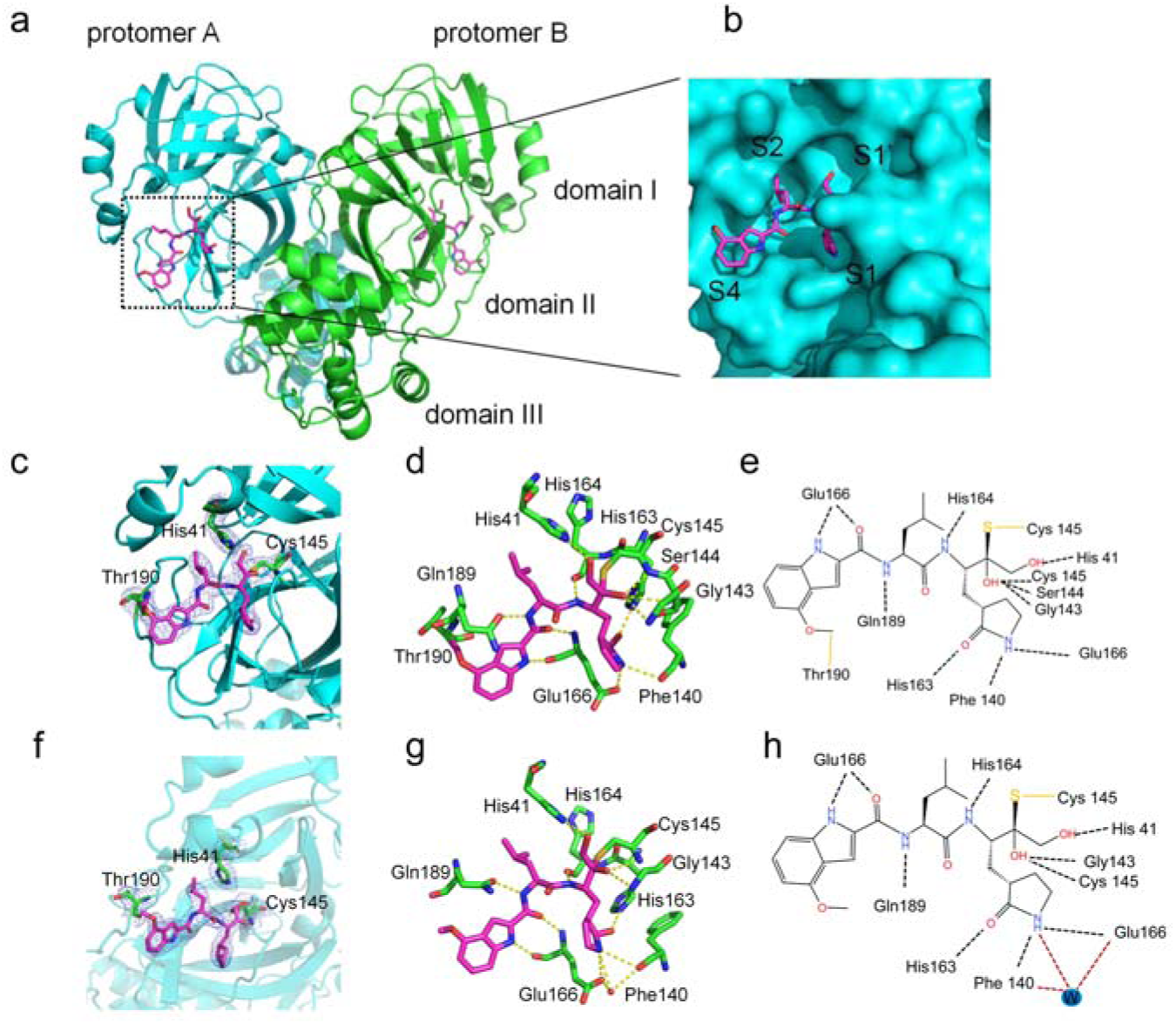
Crystal structure of SARS-CoV-2 M^pro^ in complex with PF-00835231. (a) Overall structure of SARS-CoV-2 M^pro^-PF-00835231 complex. M^pro^ is shown as cartoon. Three subdomains and two propolymers of the main protease were labeled. PF-00835231 is shown as sticks with the carbon atoms in magentas, oxygen atoms in red, nitrogen atoms in blue. (b) An enlarged view of the substrate binding pocket of SARS-CoV-2 M^pro^ with the protease shown as surface and the inhibitor shown as stick.□c to e□SARS-CoV M^pro^-PF-00835231 complex on A chain. □f to h□SARS-CoV M^pro^-PF-00835231 complex on B chain. (c and f) 2*Fo-Fc* electron density map contoured at 1.0σ.□d and e□Schematic interaction between PF-00835231 and SARS-CoV-2 M^pro^. Hydrogen-bonding interactions are indicated as dashed lines and one water molecule is labeled as W.□g and h□The detailed interaction between PF-0835231 and SARS-CoV-2 M^pro^ with residues involved in inhibitor binding (within 3.5 Å) highlighted. One water molecule is labeled W, and hydrogen bond interactions are depicted as dashed lines.

**Table 1.** Data collection and refinement statistics.

|  | MERS-5231 | SARS -5231 | SARS-Cov-2-5231 | SARS-Cov-2-G15S-5231 | SARS-Cov-2-M49I-5231 |
| --- | --- | --- | --- | --- | --- |
| **Data collection** | 8J34 | 8Z1H | 8J32 | 8J35 | 8J36 |
| Beam line | BL02U1 | BL02U1 | BL10U2 | BL10U21 | BL10U21 |
| Wavelength (Å) | 0.97918 | 0.97918 | 0.97918 | 0.97918 | 0.97918 |
| Space group | P212121 | P1 | P1211 | P1211 | P1211 |
| a, b, c (Å) | 57.90,91.33,118.14 | 55.04,60.93,68.22 | 55.17, 98.88,58.81 | 55.57,99.02,59.54 | 55.51,99.25,59.69 |
| $\alpha, \beta, \gamma$ (°) | 90., 90.,90. | 91.23,102.46,108.66 | 90.,108.00,90. | 90.,108.49,90. | 90.,108.38,90. |
| Total reflections | 264915 | 78284 | 122548 | 301714 | 416740 |
| Unique reflections | 28551 | 24344 | 28537 | 49430 | 65201 |
| Resolution (Å) | 2.30(2.43-2.30) | 2.61(2.68-2.61) | 2.21(2.33-2.21) | 1.79(1.89-1.79) | 1.65(1.74-1.65) |
| R-merge (%) | 8.5(39.8) | 7.2(13.6) | 11.3(62.3) | 2.6(68.6) | 4.4(63.1) |
| Mean I / $\sigma$ (I) | 18.5 /7.0 | 11.3 /6.0 | 8.7 /2.3 | 19.4 /2.4 | 20.6 /2.6 |
| Completeness (%) | 100.0(100.0) | 98.2(97.2) | 94.4(95.7) | 86.0(82.0) | 89.0(98.7) |
| Redundancy | 9.3(9.8) | 3.2(2.6) | 4.3(4.3) | 6.1(6.1) | 6.4(5.5) |
| **Refinement** |  |  |  |  |  |
| Resolution (Å) | 51.99-2.30 | 32.96-2.61 | 46.01-2.21 | 52.70-1.79 | 46.59-1.65 |
| Rwork/Rfree(%) | 19.82/25.55 | 18.89/23.63 | 20.44/25.42 | 20.77/24.66 | 21.19/23.44 |
| Atoms | 4675 | 4490 | 4564 | 4555 | 4616 |
| Mean temperature<br>factor (Å<sup>2</sup>) | 35.7 | 32.9 | 40.9 | 32.8 | 29.5 |
| Rmsd bond lengths<br>(Å) | 0.008 | 0.008 | 0.007 | 0.007 | 0.006 |
| Rmsd bond angles (°) | 1.118 | 1.048 | 0.973 | 0.876 | 0.904 |
| Ramachandran plot<br>(%) |  |  |  |  |  |
| Preferred | 97.81 | 97.73 | 97.13 | 98.14 | 97.63 |
| Allowed | 2.19 | 2.27 | 2.87 | 1.86 | 2.37 |
| outliers | 0 | 0 | 0 | 0 | 0 |
| Rpim | 0.029/0.133 | 0.048/0.109 | 0.060-0.326 | 0.023/0.297 | 0.019/0.298 |
| CC1/2 | 0.998/0.976 | 0.975/0.425 | 0.994/0.835 | 0.999/0.826 | 0.999/0.774 |
| Search model | 7DR8 | 7DQZ | 7C2Q | 7C2Q | 7C2Q |
| RSCC | 0.89 | 0.91 | 0.84 | 0.93 | 0.89 |
|  | 0.91 | 0.95 | 0.86 | 0.94 | 0.93 |

<sup>a</sup> The values in parentheses are for the outermost shell.
<sup>b</sup> Rfree is the Rwork based on 5% of the data excluded from the refinement.
$R_{work} = \frac{\sum |F_{obs} - F_{calc}|}{\sum F_{obs}}$ where Fobs and Fcalc are observed and calculated structure factors, respectively.

A difference was found in the binding patterns of the inhibitor with the two protomers of the dimer, and the difference mainly exists in the indole group. According to the electron density maps in both A (Figure 2c) and B protomers (Figure 2f), PF-00835231 forms an additional C-S covalent bond with the sulfur atom of Cys145. In addition, the carbon on the methoxyl group of the indole group was found to interact with Thr190. In order to further reveal the mechanism of SARS-CoV-2 M^pro^ inhibition by PF-00835231, we analyzed the interaction details between PF-00835231 and SARS-CoV-2 M^pro^ (Figure 2). The PF-00835231 employs hydroxymethyl ketone component as a directional warhead. The 4-hydroxy of PF-00835231 occupies the S1’ pocket and forms a hydrogen bond with the main chain of His41 in M^pro^. In addition, residues Gly143, His163 and Cys145 of SARS-CoV-2 M^pro^ form hydrogen bonds with the hydroxyl groups of PF-00835231. The lactam ring of PF-00835231 occupies the S1 pocket of M^pro^ and the nitrogen of lactam ring forms hydrogen bonds with Phe140, Glu166 directly or via a water molecule (W) when the carbyl oxygen of lactam forms a hydrogen bond with nitrogen of His 163. While the leucine moiety of PF-00835231 occupies the S2 pocket, The indole group of PF-00835231 occupies the S4 pocket of M^pro^. The nitrogen of the indole group and the carbyl oxygen of the main chain form a hydrogen bond with Glu166 while the nitrogen of main chain forms a hydrogen bond with His164. These can be seen on both the A (Figure 2c to 2e) and B (Figure 2f to 2h) chains, and they differ in the indole group. The carbon of the indole group methoxy is covalently bound to Thr190, which is specific to the A-chain. We found that the indole-linked methoxyl group has some interactions with Thr190, and we speculated that the oxymethyl carbon formed a covalent interaction with the carbon of Thr190. Because the electron density is not sufficient, the specific effect needs to be further verified, but it is certain that there is a certain interaction. The covalent binding of 5231 to Thr190 is present in chain A, but not in chain B, possibly because the location is dynamic. Therefore, the covalent binding to Thr190 can be seen, suggesting a novel mechanism of SARS-CoV-2 M^pro^ inhibition by PF-00835231.

Previous reports also solved the crystal structure of the SARS-CoV-2 M^pro^ in complex with PF-00835231 (PDB ID 8DSU and 6XHM) [16,26]. By superimposing the previously solved structures of SARS-CoV-2 M^pro^-PF-00835231 complex with the structure reported in this study (Figure S1), the RMSD on the 433 optimally arranged Cα atoms were 0.53 Å (PDB ID 8DSU) and 0.653 Å (PDB ID 6XHM), respectively. The binding pattern of PF-00835231 with SARS-CoV-2 M^pro^ is highly similar, except the binding mode of the indole group. The covalent binding was not found between the indole group of PF-00835231 and Thr190 of main protease in previous studies [16,26].

### Crystal structure of SARS-CoV-2 M^pro^ mutants in complex with PF-00835231

We then used the co-crystallization method to determine the crystal structures of several M^pro^ mutants of SARS-CoV-2 in complex with PF-00835231 (Figure 3). The resolutions for these structures are 1.68 Å (M^pro^ K90R with PF-00835231), 1.79 Å (M^pro^ G15S with PF-00835231), 1.91 Å (M^pro^ Y54C with PF-00835231), 1.65 Å (M^pro^ M49I with PF-00835231), 1.72 Å (M^pro^ P132H with PF-00835231), 1.64 Å (M^pro^ S46F with PF-00835231), and 1.66Å (M^pro^ V186F with PF-00835231), respectively. The data collection and detailed statistics are shown in Table 1 and Table 2. In each of these seven complex structures, each SARS-CoV-2 M^pro^ mutant molecule appears as a dimer form, which is also the form with enzymatic activity. Overall, each protomer of the SARS-CoV-2 M^pro^ mutant binds to one PF-00835231 molecule.

**Figure 3.**
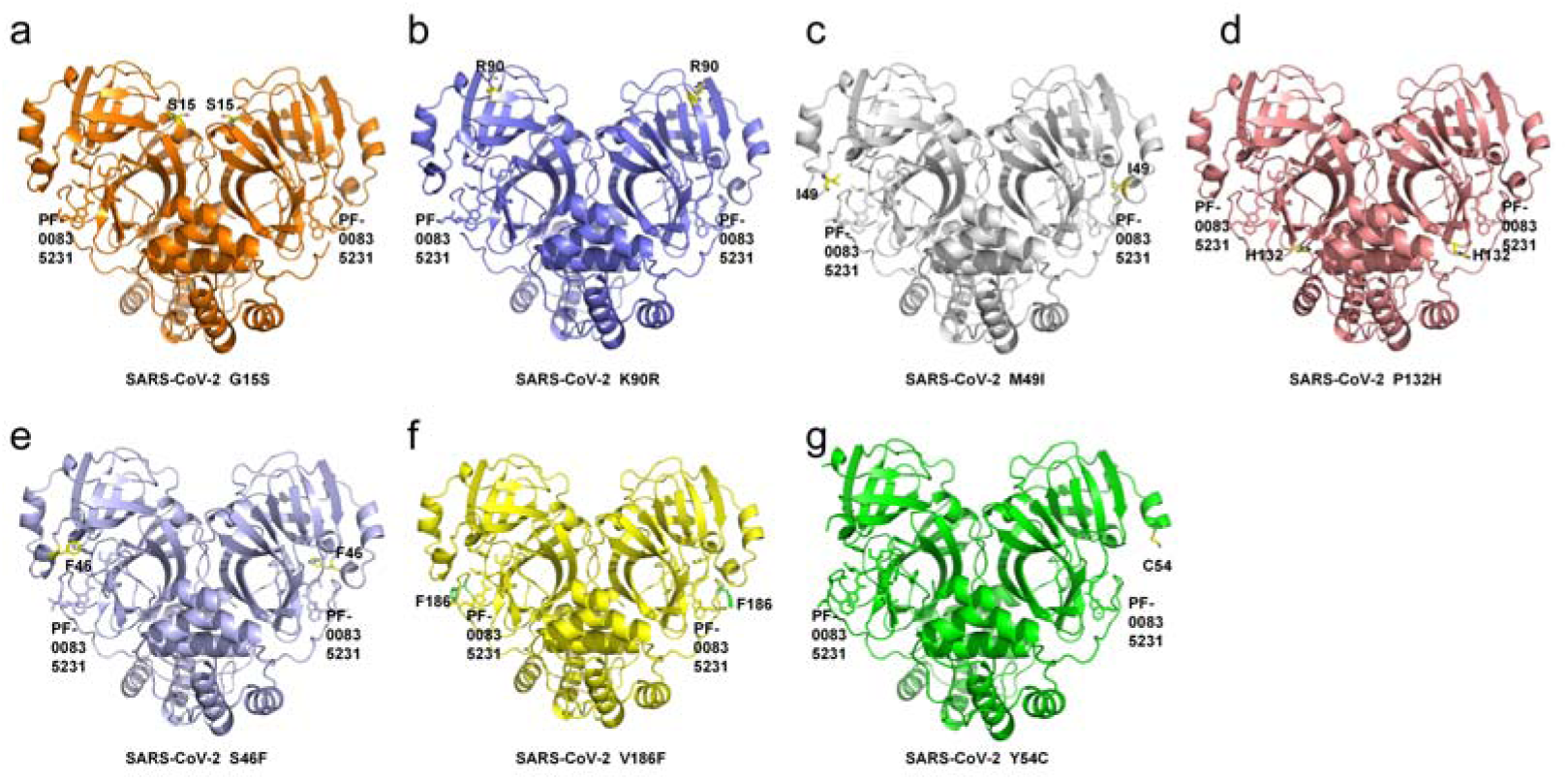
Structure overview of M^pro^ mutants from SASR-CoV-2 in complex with PF-00835231. (a) Overall structure of M^pro^ G15S-PF-00835231 complex. G15S mutant in orange cartoon while PF-00835231 is displayed in stick. (b) Overall structure of M^pro^ K90R-PF-00835231 complex. K90R mutant in slate cartoon while PF-00835231 is displayed in stick. (c) Overall structure of M^pro^ M49I-PF-00835231 complex. M49I mutant in gray90 cartoon while PF-00835231 is displayed in stick. (d) Overall structure of M^pro^ P132H-PF-00835231 complex. P132H mutant in salmon cartoon while PF-00835231 is displayed in stick. (e) Overall structure of M^pro^ S46F-PF-00835231 complex. S46F mutant in lightblue cartoon while PF-00835231 is displayed in stick. (f) Overall structure of M^pro^ V186F-PF-00835231 complex. V186F mutant in yellow cartoon while PF-00835231 is displayed in stick. (g) Overall structure of M^pro^ Y54C-PF-00835231 complex. Y54C mutant in green cartoon while PF-00835231 is displayed in stick.

**Table 2.** Data collection and refinement statistics.

|  | SARS-Cov-2-K90R-5231 | SARS-Cov-2-P132H -5231 | SARS-Cov-2-S46F-5231 | SARS-Cov-2-V186F-5231 | SARS-Cov-2-Y54C-5231 |
| --- | --- | --- | --- | --- | --- |
| **Data collection** | 8J37 | 8J38 | 8J3B | 8J39 | 8J3A |
| Beam line | BL10U2 | BL10U2 | BL10U2 | BL10U2 | BL10U2 |
| Wavelength (Å) | 0.97918 | 0.97918 | 0.97918 | 0.97918 | 0.97918 |
| Space group | P1211 | P1211 | P1211 | P1211 | P1211 |
| a, b, c (Å) | 55.63,98.98,59.56 | 55.46, 99.30,59.25 | 55.46, 99.30,59.25 | 55.45, 99.05,59.26 | 54.87, 99.93,58.71 |
| $\alpha, \beta, \gamma$ (°) | 90.,108.71,90. | 90.,108.41,90. | 90.,108.41,90. | 90.,108.37,90. | 90.,106.94,90. |
| Total reflections | 411159 | 372487 | 342391 | 400517 | 285620 |
| Unique reflections | 65079 | 57473 | 70413 | 71629 | 46688 |
| Resolution (Å) | 1.68(1.77-1.68) | 1.72(1.81-1.72) | 1.64(1.72-1.64) | 1.66(1.75-1.66) | 1.91(2.02-1.91) |
| R-merge (%) | 5.9(64.5) | 5.1(61.4) | 4.4(23.4) | 6.4(59.7) | 6.2(48.8) |
| Mean I / $\sigma$ (I) | 16.5 /2.5 | 21.1 /2.8 | 19.0 /2.8 | 14.7/2.6 | 14.4/3.0 |
| Completeness (%) | 93.8(100.0) | 88.5(99.8) | 94.2(98.8) | 99.5(100.0) | 99.8(100.0) |
| Redundancy | 6.5(5.7) | 6.5(5.9) | 4.9(2.8) | 5.6(4.9) | 6.1(5.1) |
| **Refinement** |  |  |  |  |  |
| Resolution (Å) | 46.51-1.68 | 56.21-1.72 | 48.98-1.64 | 46.41-1.66 | 37.33-1.91 |
| Rwork/Rfree(%) | 21.36/24.17 | 21.08/24.05 | 21.10/23.61 | 21.45/25.10 | 20.04/23.54 |
| Atoms | 4640 | 4623 | 4949 | 4700 | 4441 |
| Mean temperature factor (Å<sup>2</sup>) | 27.8 | 27.7 | 27.6 | 27.5 | 40.3 |
| Rmsd bond lengths (Å) | 0.006 | 0.007 | 0.007 | 0.006 | 0.007 |
| Rmsd bond angles (°) | 0.887 | 0.891 | 0.853 | 0.894 | 0.902 |
| Ramachandran plot (%) |  |  |  |  |  |
| Preferred | 97.47 | 97.47 | 97.98 | 97.64 | 97.88 |
| Allowed | 2.53 | 2.53 | 2.02 | 2.36 | 2.12 |
| outliers | 0 | 0 | 0 | 0 | 0 |
| Rpim | 0.025/0.298 | 0.022/0.276 | 0.021/0.172 | 0.030/0.301 | 0.027/0.242 |
| CC1/2 | 0.999/0.835 | 0.999/0.823 | 0.998/0.911 | 0.998/0.794 | 0.021/0.172 |
| Search model | 7C2Q | 7C2Q | 7C2Q | 7C2Q | 7C2Q |
| RSCC | 0.90 | 0.89 | 0.92 | 0.90 | 0.90 |
|  | 0.92 | 0.92 | 0.93 | 0.92 | 0.93 |

<sup>a</sup> The values in parentheses are for the outermost shell.
<sup>b</sup> Rfree is the Rwork based on 5% of the data excluded from the refinement.
$R_{work} = \frac{\sum |F_o - F_c|}{\sum F_o}$ where $F_o$ and $F_c$ are observed and calculated structure factors, respectively.

To further investigate the details of the interaction between PF-00835231 and different mutants, the amino acids that interact with PF-00835231 are clearly labeled as stick patterns (Figure 4). The interaction details between M^pro^ G15S and PF-00835231 show that PF-00835231 interacts with several residues, including His41, Cys145, His163, His164, Glu166, and Gln189 (Figure 4a). The interaction details between M^pro^ K90R and PF-00835231 show that PF-00835231 interacts with several residues, including His41, Phe140, Cys145, His163, His164, Glu166, and Gln189 (Figure 4b). The interaction details between M^pro^ M49I and PF-00835231 show that PF-00835231 interacts with several residues, including His41, Phe140, Cys145, His163, His164, Glu166, and Gln189 (Figure 4c). The interaction details between M^pro^ P132H and PF-00835231 show that PF-00835231 interacts with several residues, including His41, Phe140, Cys145, His163, His164, Glu166, and Gln189 (Figure 4d). The interaction details between M^pro^ S46F and PF-00835231 show that PF-00835231 interacts with several residues, including His41, Phe140, Cys145, His163, His164, Glu166, and Gln189 (Figure 4e). The interaction details between M^pro^ V186F and PF-00835231 show that PF-00835231 interacts with several residues, including His41, Phe140, Cys145, His163, His164, Glu166, and Gln189 (Figure 4f). The interaction details between M^pro^ Y54C and PF-00835231 show that PF-00835231 interacts with several residues, including His41, Phe140, Cys145, His163, His164, Glu166 and Gln189 (Figure 4g). Interestingly, we found in M^pro^ Y54C-PF-00835231 complex the inhibitor covalently binds to Gln189, not Thr 190, which is different from that observed in SARS-CoV-2 M^pro^-PF-00835231 complex. This further confirms the ability of PF-00835231 to form covalent bonds with the residur in S4 pocket of main protease.

**Figure 4.**
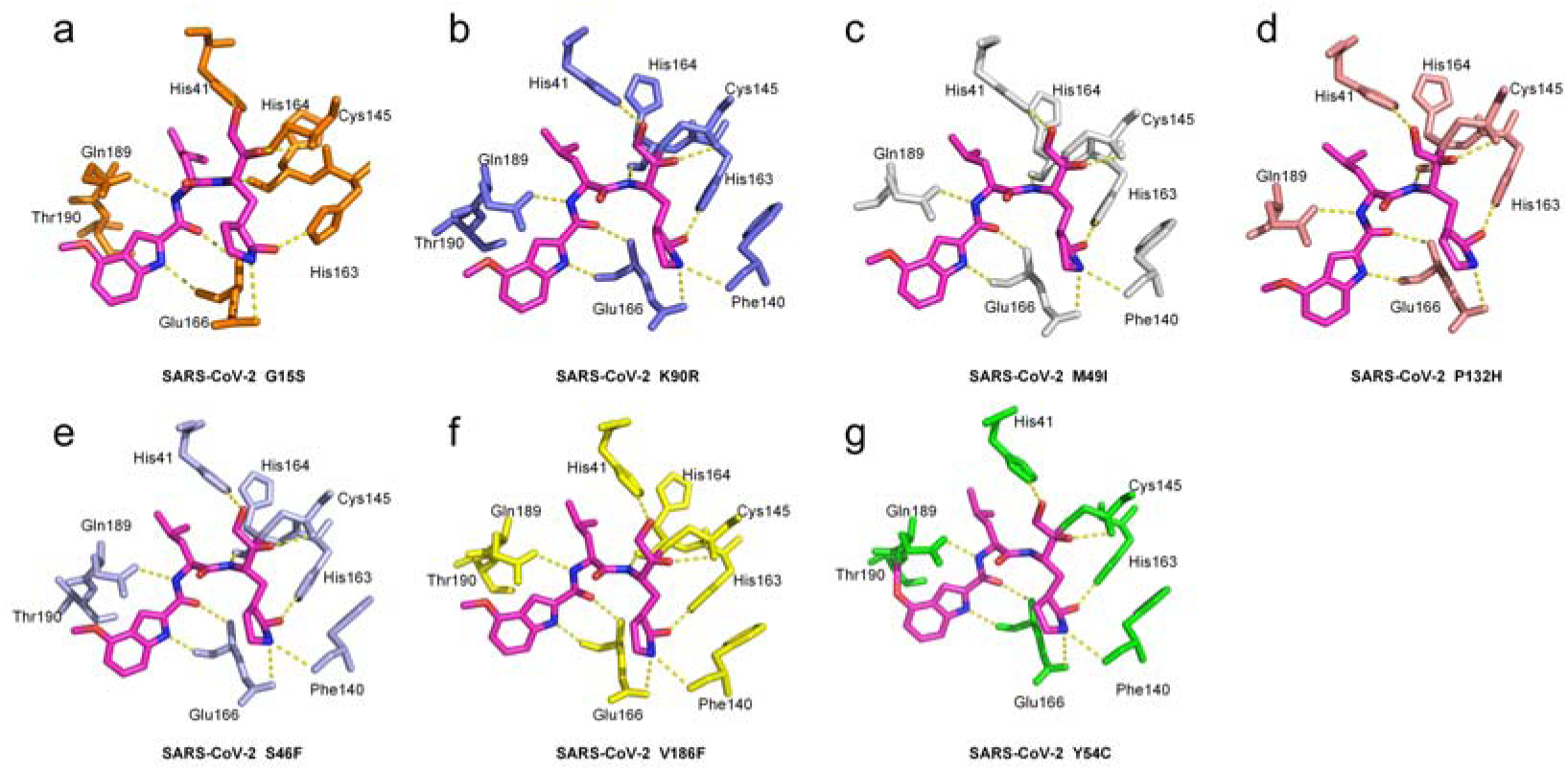
Interaction details between different M^pro^ mutants and PF-00835231. PF-00835231 is shown as sticks with the carbon atoms in magentas, oxygen atoms in red, nitrogen atoms in blue. (a) Interaction between M^pro^ G15S and PF-00835231 complex. (b) Interaction between M^pro^ K90R and PF-00835231. (c) Interaction between M^pro^ M49I and PF-00835231. (d) Interaction between M^pro^ P132H and PF-00835231. (e) Interaction between M^pro^ S46F and PF-00835231. (f) Interaction between M^pro^ V186F and PF-00835231. (g) Interaction between M^pro^ Y54C and PF-00835231.

To compare the conformational changes of M^pro^ mutants-inhibitor complexes with the wild-type M^pro^-inhibitor complex, we superimposed these structures (Figures 5, Figures S2 and Figures S3). The results clearly shows that ligand binding patterns are not disturbed by these mutations (Figures 5). The key hydrogen bond interactions between residues in wild-type M^pro^ and PF-00835231 are consistent with the structure of wild-type M^pro^-PF-00835231, except for Phe140, Gln189 and Thr 190. PF-00835231 is not found to interact with Phe140 in the M^pro^ G15S and M^pro^ P132H, and is not found to interact with Thr190 in the M^pro^ mutants. PF-00835231 is found to covalent interact with Gln189 in the M^pro^ Y54C, which is probably due to dynamic. In fact, the G15S, K90R, and P132H mutations are far from the PF-00835231 binding pocket (Figures 5b). The crystal structure revealed in this study also shows that the mutation will not cause any significant change in the inhibition potency of this inhibitor.

**Figure 5.**
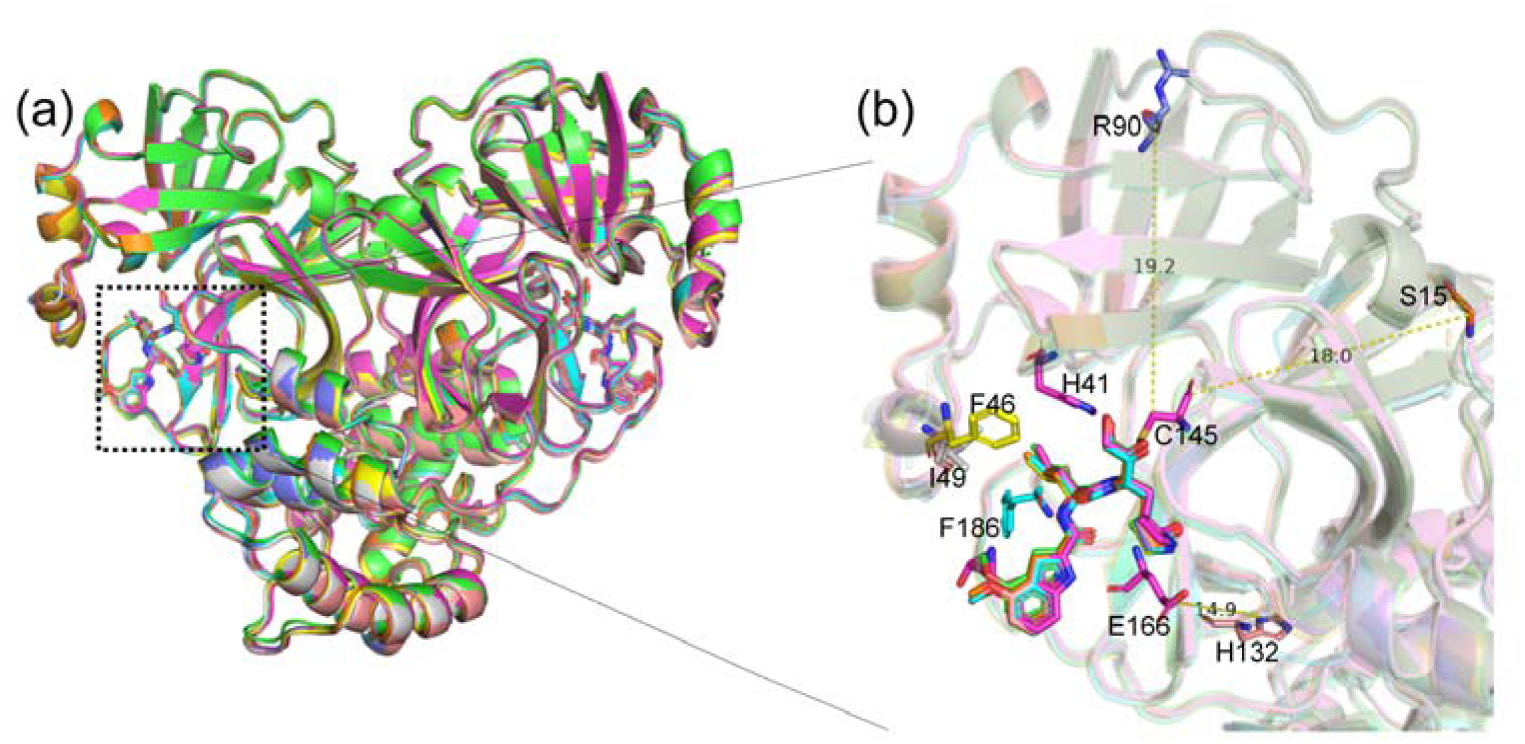
Structural comparison of PF-00835231 bound to SARS-CoV-2 M^pro^ mutants. (a)Overall structure of SARS-CoV-2 M^pro^ mutants-PF-00835231 complex. SARS-CoV-2 M^pro^-PF-00835231 is shown as cartoon with light magenta, and Y54C (green), V168F (yellow), S46F (lightblue), P132H (salmon), M49I (gray), K90R (slate), G15S (orange). (b) The zoomed-in view of structural superpositions with the location and distance of the mutant residues relative to the binding site highlighted. PF-00835231 and mutant residues are shown as sticks.

### Crystal structures of SARS-CoV and MERS-CoV M^pro^s in complex with PF-00835231

We also resolved the crystal structures of PF-00835231 in complex with SARS-CoV and MERS-CoV M^pro^s at resolutions of 2.61 Å and 2.30 Å, respectively (Table 1). We then compared the structure of SARS-CoV-2 M^pro^-PF-00835231 complex with the structures of PF-00835231 in complex with SARS-CoV and MERS-CoV M^pro^s. In terms of overall structure (Figure 6), the M^pro^s of SARS-CoV-2, SARS-CoV and MERS-CoV exhibit highly similar conformations when bound with PF-00835231. The RMSDs for the equivalent C-L positions range from 0.620 Å to 1.192 Å, respectively. As expected (Figure 7c and 7g), PF-00835231 forms a C-S covalent bond with the sulfur atom of Cys145/Cys148, same with that in SARS-CoV-2 M^pro^-PF-00835231 complex. We then analyzed the interaction details (within 3.5 Å) in SARS-CoV M^pro^-PF-00835231 complex and MERS-CoV M^pro^-PF-00835231 complex (Figure 7d and 7h). By comparison (Figure 2g and 5d), We found that the ligand-enzyme binding mode is highly similar but with some differences. The PF-00835231 directly forms hydrogen bonds with Gln89, Glu166, Phe140, His41, Ser144, Gly143, His163 in SARS-CoV M^pro^. However, the residue His164 in SARS-CoV-2 M^pro^ also forms hydrogen bonds with the methoxyketone group of PF-00835231. MERS-CoV M^pro^ binds to PF-00835231 in a highly similar way with SARS-CoV-2 M^pro^ (Figure 2g and 5g). These observations provide a structural basis for how PF-00835231 binds with M^pro^s from different coronaviruses and support that PF-00835231 can be used as a potent inhibitor with broad spectrum potential to combat diseases caused by a variety of coronaviruses.

**Figure 6.**
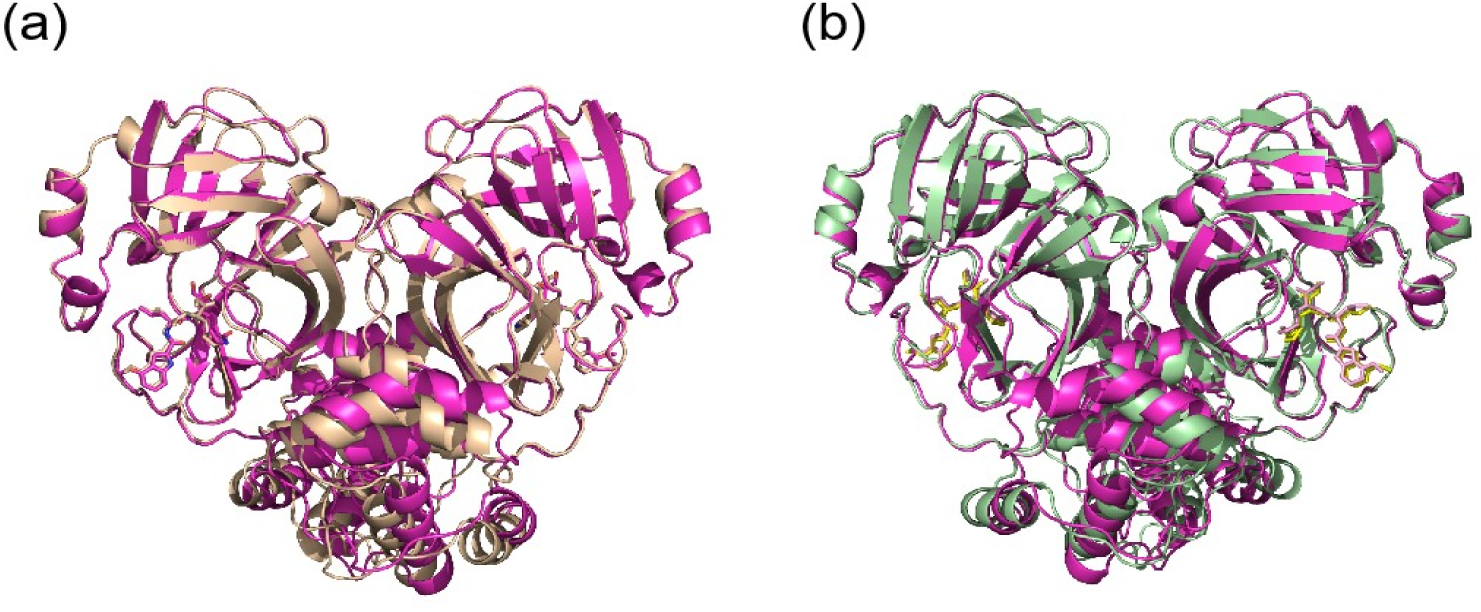
Structural comparison between different coronaviral M^pro^s in complex with PF-00835231. (a) Structural comparison between SARS-CoV-2 M^pro^-PF-00835231 (lightmagenta) and SARS-CoV M^pro^-PF-00835231(wheat). (b) Structural comparison between SARS-CoV-2 M^pro^-PF-00835231 and MERS-CoV M^pro^-PF-00835231 (palegreen).

**Figure 7.**
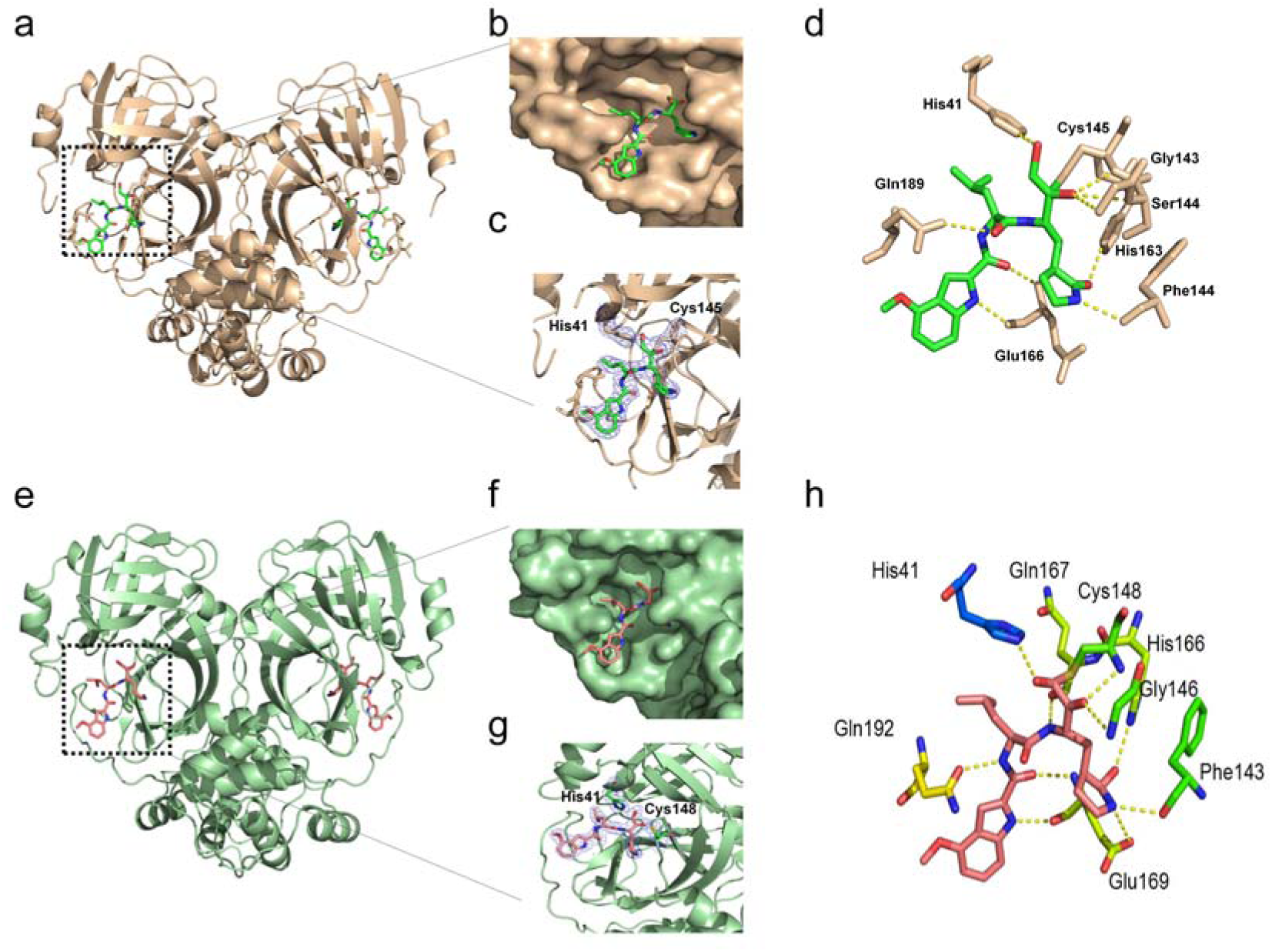
Crystal structures of SARS-CoV and MERS-CoV M^pro^s in complex with PF-00835231.(a to d) SARS-CoV M^pro^-PF-00835231 complex (wheat). (a) Overall structure of SARS-CoV M^pro^-PF-00835231 complex. M^pro^ is shown as cartoonand PF-00835231 is shown as sticks. (b) PF-00835231 in subsites of the active site of SARS-CoV M^pro^. (c) 2*Fo-Fc* electron density map contoured at 1.0σ. (d) Interaction details (within 3.5 Å) between SARS-CoV M^pro^ and PF-00835231. Hydrogen bond interactions are depicted as dashed lines. (e to h) MERS-CoV M^pro^-PF-00835231 complex (palegreen). (e) Overall structure of MERS-CoV M^pro^-PF-00835231 complex. M^pro^ is shown as cartoon. (f) PF-00835231 in subsites of the active site of MERS-CoV M^pro^. (g) 2*Fo-Fc* electron density map contoured at 1.0σ. (h) Interaction details (within 3.5 Å) between MERS-CoV M^pro^ and PF-00835231. Hydrogen bond interactions are depicted as dashed lines.

## Discussion

Over the past few years, the COVID-19 pandemic has continued to affect human health. Like other RNA viruses, SARS-CoV-2 is constantly changing through mutations, and each virus with a unique sequence is considered a new variant [27]. The continuous emergence of SARS-CoV-2 variant has seriously endangered public health security. Continued research on these inhibitors is conducive to a rapid understanding of their current and future antiviral effects. Among several structural and non-structural SARS-CoV-2 proteins, M^pro^ has been designated as a potential therapeutic target for drug development [13,15]. Inhibiting M^pro^ would prevent virus replication and constitute one of the potential anti-coronavirus strategies. The development of anti-coronavirus M^pro^ inhibitors has aroused great interest. A recombinant compound that is the hydroxymethyl ketone covalent inhibitor PF-00835231 has entered clinical trials [16]. This phosphate prodrug, PF-07304814, is converted to its active form, PF-00835231, and irreversibly attaches to the active Cys. Here,we evaluate the details of interactions between the small hydroxymethylketone-based PF-00835231 molecule and seven different M^pro^ mutants.

There is an interesting finding in our study --the carbon on the methoxyl group of the indole group of PF-00835231 was found to interact with Thr190 in SARS-CoV-2 M^pro^, while interacts with Gln189 in M^pro^ Y54C. However, we didn’t find that it interacts with any residues in other SARS-CoV-2 M^pro^ mutants and SARS-CoV M^pro^ and MERS-CoV M^pro^s possibly because of dynamic in the location. Though the electron density is not high enough, the specific effect needs to be further verified, but it is certain that there is a certain interaction. This suggests some new possibilities for the mechanism of SARS-CoV-2 M^pro^ inhibition by PF-00835231.

In this study, we solved the crystal structure of PF-00835231 and SARS-CoV-2 M^pro^ complexes, and also resolved the crystal structure of PF-00835231 with SARS-CoV and MERS-CoV M^pro^s complexes. These structures suggest that PF-00835231 has similar binding patterns for different M^pro^s, but with subtle differences. These data update previous reports on the discovery of PF-00835231 and contribute to a comprehensive understanding of the inhibition mechanism of PF-00835231. Our study also determined the crystal structures of PF-00835231 in complex with of several SARS-CoV-2 M^pro^ mutants. Our data shown that the binding mode of PF-00835231 is not significantly affected by these mutants. The binding pattern of PF-00835231 with SARS-CoV-2 M^pro^ mutants, SARS-CoV M^pro^ and MERS-CoV M^pro^s and SARS-CoV-2 variants is similar.

Because M^pro^ is highly conserved in different human coronaviruses, potent inhibitors like PF-00835231 can also be used as broad-spectrum candidates for a variety of coronavirus infections. Therefore, this study provides a theoretical basis for the treatment of SARS-CoV-2 M^pro^ mutants, SARS-CoV M^pro^ and MERS-CoV M^pro^s and SARS-CoV-2 variants and further drug development.

---

**Figure S1.**
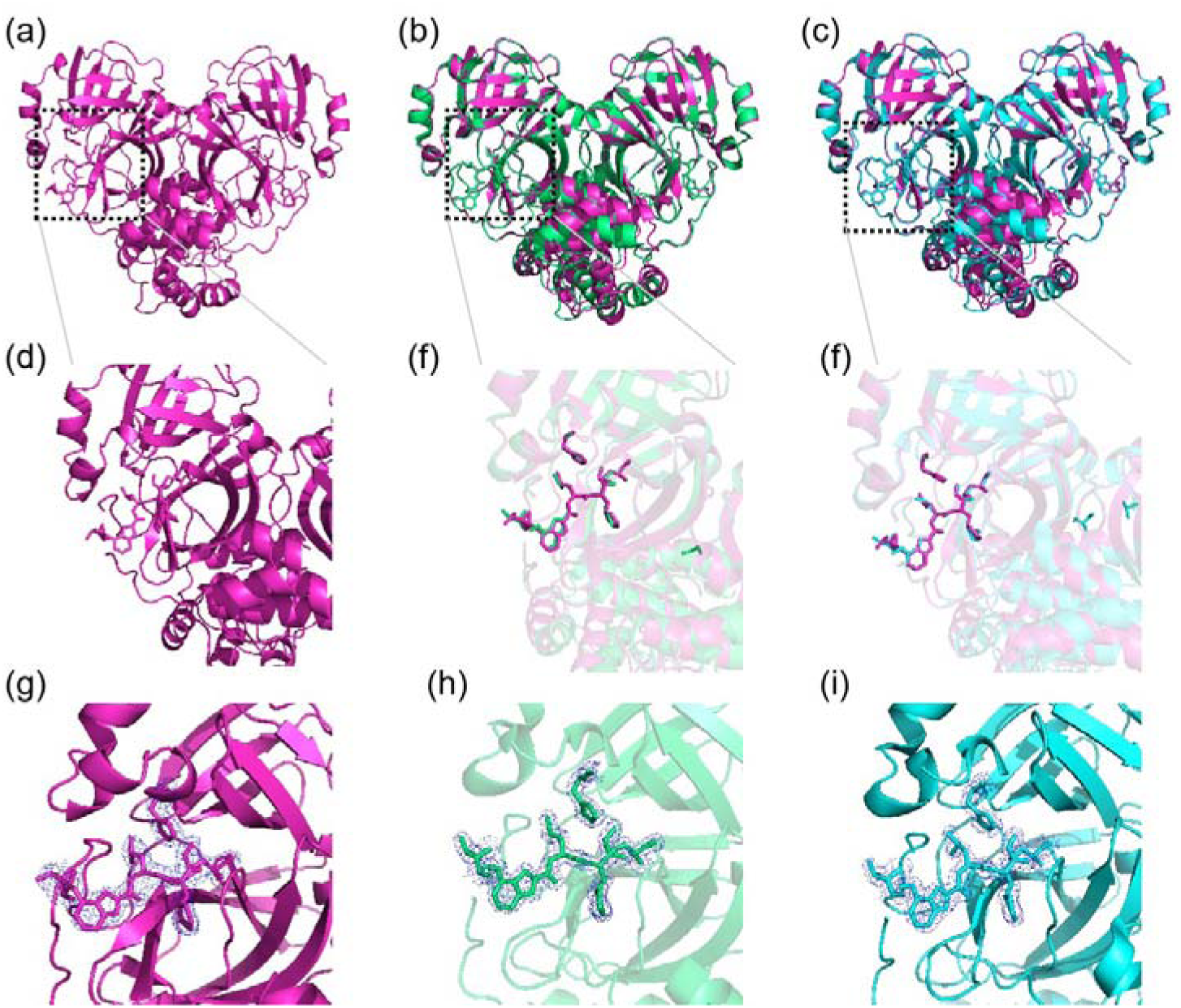
Structural comparison between SARS-CoV-2 M^pro^-PF-00835231 and available structures. (a)Structural of SARS-CoV-2 M^pro^-PF-00835231.(b)Structural comparison between SARS-CoV-2 M^pro^-PF-00835231 and SARS-CoV-2 M^pro^-PF-00835231 (PDB ID 6XHM, limegreen). (c) Structural comparison between SARS-CoV-2 M^pro^-PF-00835231 and SARS-CoV-2 M^pro^-PF-00835231 (PDB ID 8DSU, cyan). SARS-CoV-2 M^pro^-PF-00835231 is shown as cartoon with light magenta (thisresearch), limegreen (PDB ID 6XHM) and cyan (PDB ID 8DSU).□d-f□The zoomed-in view of the substrate binding pocket of main proteases. PF-00835231 and catalytic dyad residues are shown as sticks. (g) The 2*Fo-Fc* electron density map (contoured at 1.0σ) of the inhibitor and catalytic dyad residues, His41,Cys145 and Thr190, in SARS-CoV-2 M^pro^-PF-00835231 complex (this study) is shown as blue mesh. (h) The 2*Fo-Fc* electron density map (contoured at 1.0σ) of the inhibitor and catalytic dyad residues, His41, Cys145 and Thr190, in SARS-CoV-2 M^pro^-PF-00835231 complex (PDB ID 6XHM) is shown as blue mesh. (i) The 2*Fo-Fc* electron density map (contoured at 1.0σ) of the inhibitor and catalytic dyad residues, His41,Cys145 and Thr190, in SARS-CoV-2 M^pro^-PF-00835231 complex (PDB ID 8DSU) is shown as blue mesh .

**Figure S2.**
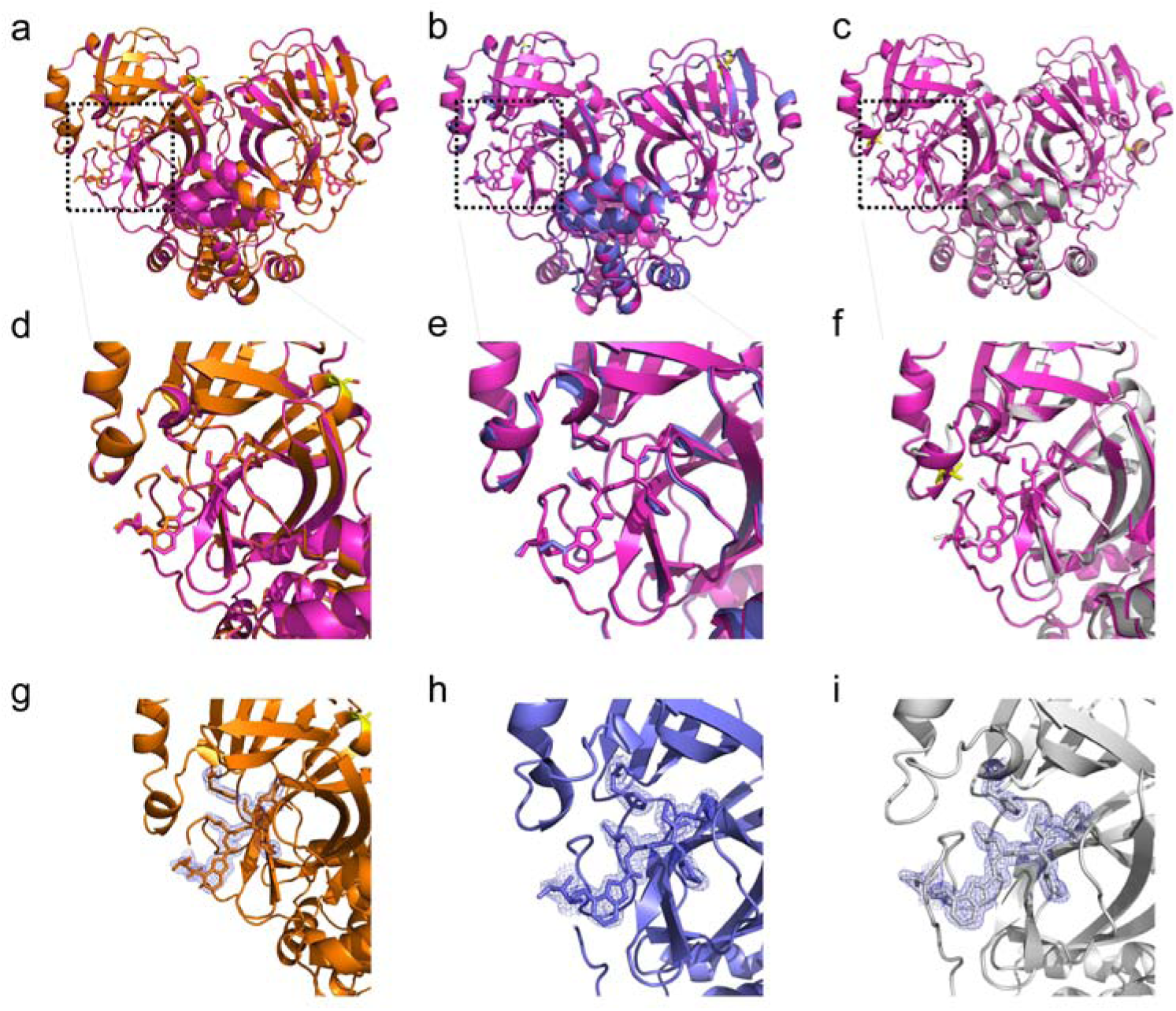
Structural comparison between SARS-CoV-2 M^pro^-PF-00835231 and M^pro^ mutants-PF-00835231 (G15S, K90R and M49I).The S15,R90 and I49 are show as yellow sticks. (a) Structural comparison between SARS-CoV-2 M^pro^-PF-00835231 and G15S-PF-00835231 (b) Structural comparison between SARS-CoV-2 M^pro^-PF-00835231 and M^pro^ K90R-PF-00835231. (c) Structural comparison between SARS-CoV-2 M^pro^-PF-00835231 and M^pro^ M49I-PF-00835231.□d-f□The zoomed-in view of the substrate binding pocket of M^pro^ and M^pro^ G15S. PF-00835231 and catalytic dyad residues are shown as sticks. (g-h) The 2*Fo-Fc* electron density maps of the PF-00835231 bound to different SARS-CoV-2 M^pro^ mutants. 2*Fo-Fc* electron density maps (blue) were contoured at 1 σ. (g) The 2*Fo-Fc* electron density map of PF-00835231 bound to G15S mutant. (h) The 2*Fo-Fc* electron density map of PF-00835231 bound to K90R mutant. (i) The 2*Fo-Fc* electron density map of PF-00835231 bound to M49I mutant.

**Figure S3.**
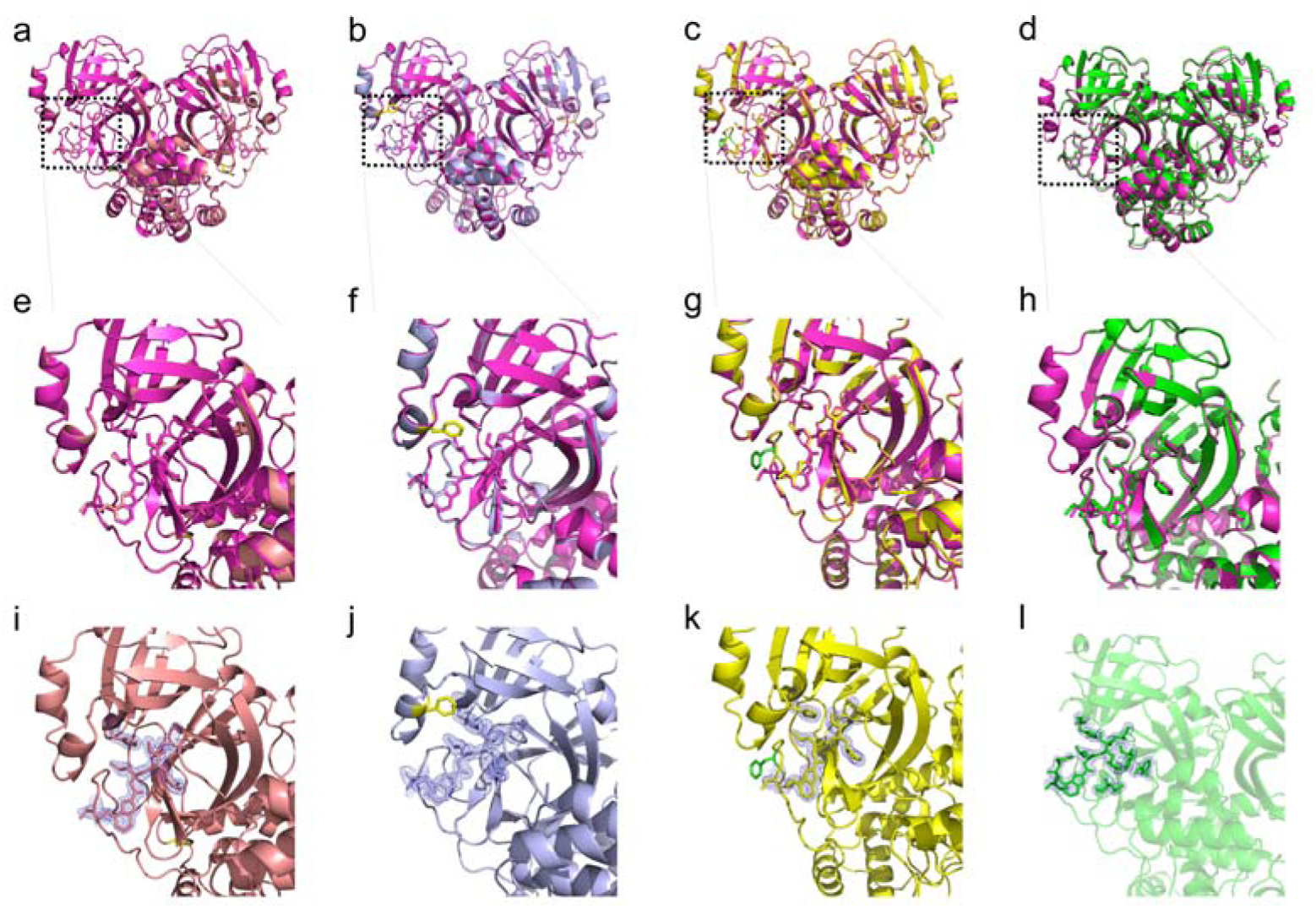
Structural comparison between SARS-CoV-2 M^pro^-PF-00835231 and M^pro^ mutants-PF-00835231 (P132H, S46F,V186F and Y54C). The H132, F46 and C54 are show as yellow sticks while F186 is shou as green stick. (a) Structural comparison between SARS-CoV-2 M^pro^-PF-00835231 and P132H-PF-00835231 (b) Structural comparison between SARS-CoV-2 M^pro^-PF-00835231 and M^pro^ S46F-PF-00835231. (c) Structural comparison between SARS-CoV-2 M^pro^-PF-00835231 and M^pro^ V186F-PF-00835231. (d)Structural comparison between SARS-CoV-2 M^pro^-PF-00835231 and M^pro^ Y54C-PF-00835231.□e-h□The zoomed-in view of the substrate binding pocket of M^pro^ and M^pro^ G15S. PF-00835231 and catalytic dyad residues are shown as sticks. (i-l) The 2*Fo-Fc* electron density maps of the PF-00835231 bound to different SARS-CoV-2 M^pro^ mutants. 2*Fo-Fc* electron density maps (blue) were contoured at 1 σ. (i) The 2*Fo-Fc* electron density map of PF-00835231 bound to P132H mutant. (j) The 2*Fo-Fc* electron density map of PF-00835231 bound to S46F mutant. (k) The 2*Fo-Fc* electron density map of PF-00835231 bound to V186F mutant.(l) The 2*Fo-Fc* electron density map of PF-00835231 bound to Y54C mutant.

